# HSC70, HSPA1A, and HSP90AB1 Facilitate Ebola Virus trVLPs to Induce Autophagy

**DOI:** 10.1101/2020.06.22.163683

**Authors:** Dong-Shan Yu, Shu-Hao Yao, Wen-Na Xi, Lin-Fang Cheng, Fu-Min Liu, Hai-Bo Wu, Xiang-Yun Lu, Nan-Ping Wu, Shui-Lin Sun, Hang-Ping Yao

## Abstract

Ebola virus (EBOV) can induce autophagy to benefit the virus life cycle, but detailed mechanisms remain to be elucidated. We previously found that EBOV GP and VP40 proteins interact with HSC70 (HSPA8), HSPA1A, and HSP90AB1. Thus, we presumed that EBOV likely induced autophagy by virus protein-host HSC70 or co-chaperon interactions via chaperone-mediated autophagy (CMA). We developed EBOV-trVLPs to model the EBOV life cycle, infected 293T cells with trVLPs, evaluated CMA by GFP-LC3 and RFP-LAMP1 co-localization, transmission electron microscopy (TEM) observation, and immunoblot analysis. The results demonstrated that EBOV-trVLPs induce autophagy which could not be inhibited by 3-MA significantly; autophagosomes and autolysosomes are obviously in the cytoplasm confirming CMA in cells infected with trVLPs. Meanwhile, a knockdown of HSC70 and relevant co-chaperones could inhibit trVLPs-associated autophagy, but no effort to Akt/mTOR/PHLPP1 pathway. These data indicate that EBOV-trVLPs could induce autophagy by CMA but was not limited by the CMA pathway. HSC70, HSPA1A, and HSP90AB1 participate and regulate CMA induced by EBOV-trVLPs. This was the first study about EBOV-trVLPs-induction of CMA and provides insight into the viral protein-host protein interaction, which is probably associated with CMA.

**Highlights:**

- EBOV-trVLPs induce chaperone-mediated autophagy (CMA) but are not limit by CMA.
- HSC70, HSPA1A, and HSP90AB1 facilitate EBOV-trVLPs to induce autophagy
- Knockdown of HSC70, HSPA1A, and HSP90AB1 inhibit EBOV-trVLPs-induced CMA.

## INTRODUCTION

Ebola virus (EBOV) was first reported in 1976 as the pathogen of the highly contagious zoonotic disease and Ebola viral disease (EVD) both in humans and other primates (Emond et al., 1977). EBOV is a single-stranded, negative-sense RNA virus belongs to the filovirus family, according the epidemic focus, five different EBOVs have been defined: 1) Ebola virus (EBOV, previously known as Zaire ebolavirus); 2) Sudan virus (SUDV); 3) Bundibugyo virus (BDBV); 4) Taï Forest virus (TAFV); and 5) Reston virus (RESTV) (Kuhn, 2017). From December 2013 to March 2016, EBOV caused EVD outbreak in Western Africa with more than 28,000 cases and 11,000 deaths in 11 countries(Agua-Agum et al., 2016). In 2019, the ongoing EVD epidemic was significantly expanded and was associated with a high fatality rate of 67% from 2018 to 2019 in the democratic public of the Congo (Ilunga Kalenga et al., 2019).

Although significant progresses have been devoted to EVD therapeutics, the pathogenesis, molecular mechanisms, and EBOV-host interactions remain poorly understood. EBOV-induced autophagy is an area in which the associated mechanisms remain unclear. Autophagy is a regulated process responsible for degradation and recycling of damaged organelles through lysosomal degradation (Jackson, 2015). Under conditions of cellular stress (e.g., nutrient deficiency, virus infection, and hemotherapy), autophagy can be initiated through different mechanisms and affect the survival of virally infected or transformed cells (Levine and Kroemer, 2008; Choi et al., 2013; Parzych and Klionsky, 2014). There are at least three types of autophagy have been found in mammalian cells: 1) macroautophagy; 2) microautophagy; and 3) chaperone-mediated autophagy (CMA) (Madrigal-Matute and Cuervo, 2016). CMA is a selective degradation pathway with cargo recognition and internalization into the lysosome. CMA start with a recognizing of cytosolic heat shock cognate protein of 70 kDa (HSC70) to a consensus pentapeptide motif (KFERQ-like motif) (Sahu et al., 2011; Kaushik and Cuervo, 2012). A group of co-chaperones, including HSP90, BAG-1, and Hip then form a complex of HSC70/KFERQ-containing proteins. The HSC70/KFERQ-protein complex targets the lysosome by binding to lysosome-associated membrane protein type 2A (LAMP-2A), which is recognized as an HSC70 receptor, and is then transported through the membrane and is degraded in the lysosomal lumen (Chiang et al., 1989; Bandyopadhyay and Cuervo, 2008; Bandyopadhyay et al., 2008).

Currently, three kinds of mechanisms are possibly involved in virus-induced autophagy: 1) double stranded RNA facilitates autophagosome by down-regulation of mTOR and PKR activity; 2) virus infection induces autophagosome formation through ER stress; and 3) virus infection activates mTOR and accelerates cellular translation (Kroemer et al., 2010; Sancak et al., 2010; Kaushik and Cuervo, 2012). To date, the connection between EBOV and autophagy remains poorly understood. It has been reported that the BCL2-associated athanogene-3 (BAG-3) WW-domain can interact with the PPxY motif of both EBOV and MARV VP40, which negatively regulates the budding of VP40 VLPs and infectious virus (Liang et al., 2017). Previous studies also indicate that a knockout of the endoplasmic reticulum (ER)-resident protein, FAM134B, resulted in an inhibition of replication of the Ebola virus Strains, Makona and Mayinga (Chiramel et al., 2016). However, the detailed mechanisms of how EBOV induces autophagy remain to be elucidated.

In our previous studies, we found that the EBOV GP protein interacts with HSC70 (HSPA8), HSPA1A, and HSP90AB1; the EBOV VP40 protein and nucleic acids both interact with HSPA1A and HSP90AB1(Yu et al., 2018). Based on previous research, we presumed that EBOV could most likely induce autophagy by virus protein-host HSC70 or co-chaperon interaction via CMA in some yet-undefined pathways. Thus, in the present study, we used plasmids to co-transfected 293T cells to produce transcription- and replication-competent virus-like particles (trVLPs) in order to model the EBOV life cycle under biosafety level 2 conditions (Hoenen et al., 2014), assessed autophagy induced by trVLPs, and evaluated CMA based on the co-localization of microtubule-associated protein light chain 3 (LC3) and lysosome-associated membrane glycoprotein 1 (LAMP1). We also used transmission electron microscopy (TEM) observations, knockdown of host factors, and immunoblot analysis to assess EBOV-trVLPs and host elements involved in the autophagy and CMA pathways. This is the first survey of the EBOV-trVLPs life cycle and CMA pathway.

## MATERIALS AND METHODS

### Cell Lines and Plasmids

Human embryonic kidney (HEK) 293T cells were maintained in Dulbecco’s modified Eagle’s medium (DMEM; Thermo Fisher, Waltham, MA, USA; Cat#10566016) supplemented with 10% fetal calf serum (FBS; Gibco, Waltham, MA, USA; Cat#10099141), 2 mM L-glutamine (Life Technologies, Waltham, MA, USA; Cat#25030081), and 1% penicillin-streptomycin (Life Technologies; Cat#10378016) at 37°C in a humidified 5% CO2 incubator with 5% CO2. Plasmids pCAGGS-L, pCAGGS-NP, p4cis-vRNA-RLuc, pCAGGS-Tim1, pCAGGS-T7, pCAGGS-VP30 and pCAGGS-VP35 were kindly provided by Drs. Heinz Feldmann and Thomas Hoenen, Rocky Mountain Laboratories, National Institute of Health. GFP-LC3 (Addgene, Watertown, MA, USA; Cat# 11546) and RFP-LAMP1 (Addgene, Cat# 1817) were purchased from Addgene company.

### Production of trVLPs

Plasmids (75 ng pCAGGS-VP30, 125 ng pCAGGS-VP35, 125 ng pCAGGS-NP, 1,000 ng pCAGGS-L, 250 ng p4cis-vRNA-RLuc and 250 ng pCAGGS-T7) were use d to co-transfected producer 293T cells (p0 generation) to produce trVLPs (Mizushima et al., 2010). Cellular supernatants containing released trVLPs were harvested at 72 h post-transfection and used to infect target 293T cells (p1 generation) which previously transfected with plasmids (125 ng pCAGGS-NP, 125 ng pCAGGS-VP35, 75 ng pCAGGS-VP30, 1,000 ng pCAGGS-L, and 250 ng pCAGGS-Tim1) (Hoenen et al., 2014). 72 h after transfection, the supernatants were centrifuged at 175 ×g to get rid of cell debris. The cleared supernatants in 2 mL of 20% sucrose from the bottom of each tube were centrifuged at 125,000 × g in a SW-28 rotor for 3 h at 4°C. The resulting pellets were resuspended in 100 μL ice-cold NTE buffer (10 mM Tris pH 7.5, 100 mM NaCl, and 1 mM EDTA) by tapping gently approximately 100 times, and trVLPs were stored on ice or in a refrigerator at 4°C until further use.

### TCID50 and MOI of trVLPs

TrVLPs infectivity titre was titrated using 50% tissue culture infective dose (TCID50). TrVLPs were diluted at 10-fold serially with DMEM from 10^−1^ to 10^−10^ concentrations. The attenuated trVLPs (100 μl) were added to wells in each row of a 96-well plate, and 293T cell suspension (100 μl) was added to each well to a final cell density of 2×10^5^ cells/ml. 293T cells without HP001 infection were included as controls. Cytopathic effects (CPEs) were observed by microscopy and recorded each day for 7 days, the results were used to calculate TCID50 by the Reed-Muench method(Ramakrishnan, 2016). The MOI value was calculated according the equation MOI=0.7×TCID50/ cell numbers (Supplementary Table 1).

### Electron Microscopy (EM) Analysis of trVLPs

Fresh trVLPs (10μl) were pipetted on a 300-mesh copper grids, 5 min after incubation, the grids were washed twice with distilled water and stained with 1% uranyl acetate for 15 sec. Then, the excess liquid were removed with filter paper, the trVLPs were watched under transmission electron microscope (H-7000, Hitachi, Hitachinaka, Japan).

### Immunofluorescence

The 293T cells cultured in 24-wells plates were transfected with plasmids GFP-LC3 (800 ng/well), or GFP-LC3 plus LAMP1-RFP (400 ng/well each). 24 h after transfection, the cells were treated with trVLPs (MOI = 1.5), or rapamycin (0.5 nM), or trVLPs (MOI = 1.5) plus 3-MA (0.5 nM). At 48 h after trVLPs infection or drug treatment, the GFP-LC3 and LAMP1-RFP dot formations were detected under a fluorescence microscope (DMi8-M, Leica, German). The cell containing ≥ 5 GFP-LC3 dots is defined as autophagy positive cells (Mizushima et al., 2010).

### TEM Analysis of Autophagy

48 h after trVLPs (MOI = 1.5) infection or 3-MA (0.5 nM) treatment, 293T cells were collected and fixed with 2.5% glutaraldehyde overnight at 4°C. Later, the cells were treated with 0.1% OsO4 solution for 2 h, then dehydrated in a graded series of ethanol and embedded in epoxy resin 812. Ultrathin sections of specimens were collected on copper grids, double-stained with uranyl acetate and lead citrate, and examined by transmission electron microscopy (H-7000, Hitachi, Hitachinaka, Japan).

### Western blot analysis

Total protein from 293T cells after treatment was extracted by extraction buffer RIPA (Solarbio, Beijing, China, Cat# R0010) with protease inhibitor PMSF (Solarbio, Cat# P6730). Protein concentration was quantified with the bicinchoninic acid (BCA) method (Sigma, Santa Clara, CA, USA, Cat#BCA1-1KT). The extract was eluted with 2× sodium dodecyl sulfate (SDS) Loading Buffer and resolved by SDS-polyacrylamide gel electrophoresis (PAGE). Proteins were transferred to polyvinylidene fluoride (PVDF) membranes, incubated with the relevant primary and secondary antibodies, detected by the Super Signal West Pico chemiluminescent substrate (Thermo Fisher; Cat# 34580). GAPDH was used as internal control. When immunoblot analysis was performed with lysosomal samples, LAMP2 protein was used as internal control.

### Antibodies

Antibodies recognizing LC3B (Cat# 3868S), Phospho-mTOR(Cat# 5536S), Phospho-Akt (Cat# 4060S), HSPA1A (HSP70; Cat# 4873S), LAMP1 (Cat# 15665s), LAMP2 (Cat# 49067S), GAPDH (Cat# 2188), HRP-linked anti-rabbit IgG (Cat# 7074S), and HRP-linked anti-mouse IgG (Cat# 7076S) were purchased from Cell Signalling Technology (Danvers, MA, USA). Rapamycin (Cat# 1912) and 3-Methyladenine (3-MA, Cat# 3977) were purchased from R&D Systems (Minneapolis, MN, USA). HSC70 (HSPA8, Cat# NBP2-12880) and HSP90AB1 (Cat# NB110-61640) were purchased from Novus Biologicals (Littleton, CO, USA). PHLPP1 (Cat# MBS151247) was obtained from MyBioSource company (San Diego, CA, USA).

### SiRNA knockdown

SiRNAs targeting HSC70, HSPA1A and HSP90AB1 were designed and synthesized with three siRNA sequences for each one, the most efficient sequence for RNA interference (RNAi) was evaluated by qPCR and were chosen for tests (Supplementary Table 2). Meanwhile, cytotoxity of siRNAs were estimated by nuclei counting, the siRNA was classified as hypotoxicity if 75 or more nuclei within one vision of microscope (40×, 0.24mm^2^) (Supplementary Figure 1). In the experiments, 100 μl of opti-MEM medium (Invitrogen; Cat# 31985070) containing 1.4 μl siRNA and 4.5 μl HiPerFect transfection reagent (Qiagen, Dusseldorf, Germany; Cat# 301705) was added to each well of 24-well plates, plates were shaken gently for 1 min. After 10 min incubation at room temperature, a cell suspension (400 μl) containing 1×10^5^ cells were added to give a final siRNA concentration of 75 nM. Cells were incubated at 37°C and 5% CO2 for 48 h.

### Lysosome Isolation

Lysosomes were purified from 293T cells using a lysosome isolation kit (Sigma, Cat# LYSISO1-1KT). According the protocol, 1 × 10^8^ 293T cells were collected and the cell membranes were disrupted by ultrasound. The breakage was evaluated by staining with Trypan blue. The specimen was then serially centrifuged at 1,000 × g and 20,000 × g. The pellet was collected, and calcium chloride was later added to 8 mM, incubated for 15 min at room temperature, centrifuged at 5,000 × g, and the supernatants that contained lysosomal matrices were collected.

### Statistical Analysis

Data were presented as the mean ± standard deviation in all quantitative experiments. Student’s t-tests (two-tailed, or paired t-tests) were performed where appropriate. All analysis was performed with Graphpad Prism 8 (GraphPad Software, UK). Differences between groups with p <0.05 were considered statistically significant.

## RESULTS

### EBOV-trVLPs could strongly mimic the wild type Ebola virus strain

EBOV-trVLPs exhibited a filamentous-like morphology with a major axis that was approximately 100 nm − 1000 nm and a diameter of about 50 nm as visualized by EM (Fig. 1A). Meanwhile, DiI-labeled trVLPs in live 293T cells were visualized by fluorescent microscopy which indicated trVLPs could translocated into the cytoplasm (Fig. 1B). The images indicated that trVLPs could simulate the wild type strain in morphology, size, and basic functions to the greatest extent (Nanbo et al., 2010).

**Figure 1.**
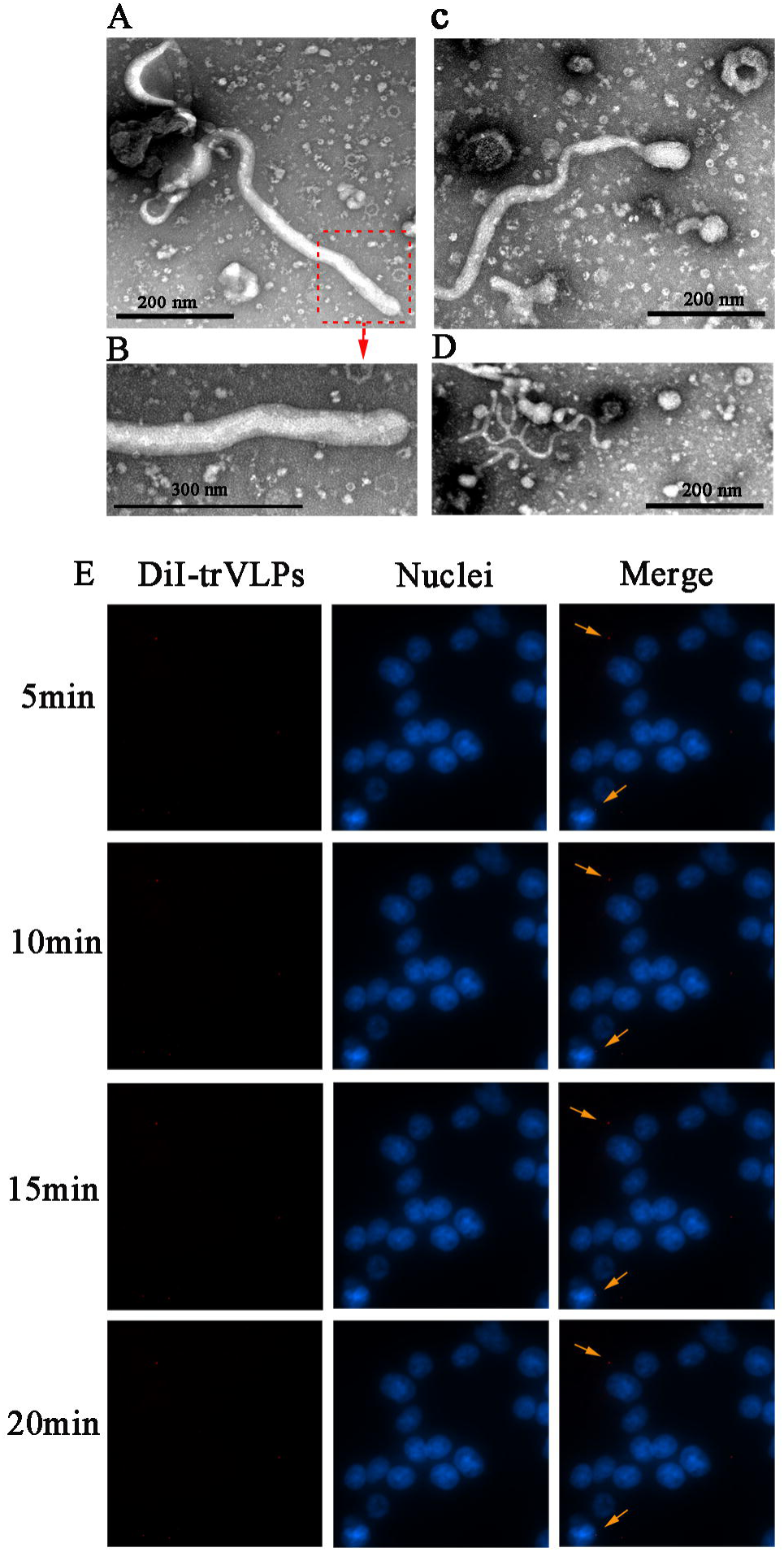
Production of EBOV-trVLPs. **(A)** Visualization of EBOV-trVLPs by EM. trVLPs were produced according to the protocol, purified by ultracentrifugation using a 20% sucrose gradient, and virion particles were visualized by EM following negative staining. The trVLPs exhibit filamentous-like viral particles. **(B)** Cellular internalization of DiI-labeled trVLPs. trVLPs were added to DiI label, which could be traced by fluorescence microscopy. At 5, 10, 15, and 20 min after 293T cells were infected, the DiI-trVLPs shows the activity of the invading cells (indicated by yellow arrows). EBOV-trVLPs, ebolavirus transcription- and replication-competent virus-like particles; EM, electron microscopy; DiI, 1,1’-dioctadecyl-3,3,3’,3’-tetramethyl indocarbocyanine perchlorate.

### EBOV-trVLPs induced autophagy in 293T cells

At 48 h after infection with trVLPs, autophagy was observed in 293T cells. As Fig. 2A indicates, the cells containing GFP-LC3 punctate structures used to monitor the number of autophagosomes were dominant among the GFP-LC3-transfected cells. At a high magnification, the punctate dots were rich in these cells (≥ 5 dots/cell) (Fig. 2B and C). Moreover, we investigated the GFP-LC3-I and GFP-LC3-II proteins extracted from the cells by immunoblotting with antibodies against LC3 (Fig. 2D). The results indicated that the cells induced by rapamycin or infected by trVLPs exhibited significant GFP-LC3-II expression, and the conversion from GFP-LC3-I to GFPLC3-II indicated the process of autophagy. Cells previously treated with 3-MA and subsequently infected with trVLPs also displayed substantial GFP-LC3-II expression, which indicated that 3-MA could not effectively inhibit autophagy induced by trVLPs. 3-MA is one of the most commonly used pharmacological approaches to inhibit class I PI3-kinase (PI3-K) and class III PI3 kinase activity which is required for autophagy (Liang et al., 2016). The results indicate that trVLPs probably induced autophagy through other means and was not limited to the PI3-K/AKT/mTOR pathway.

**Figure 2.**
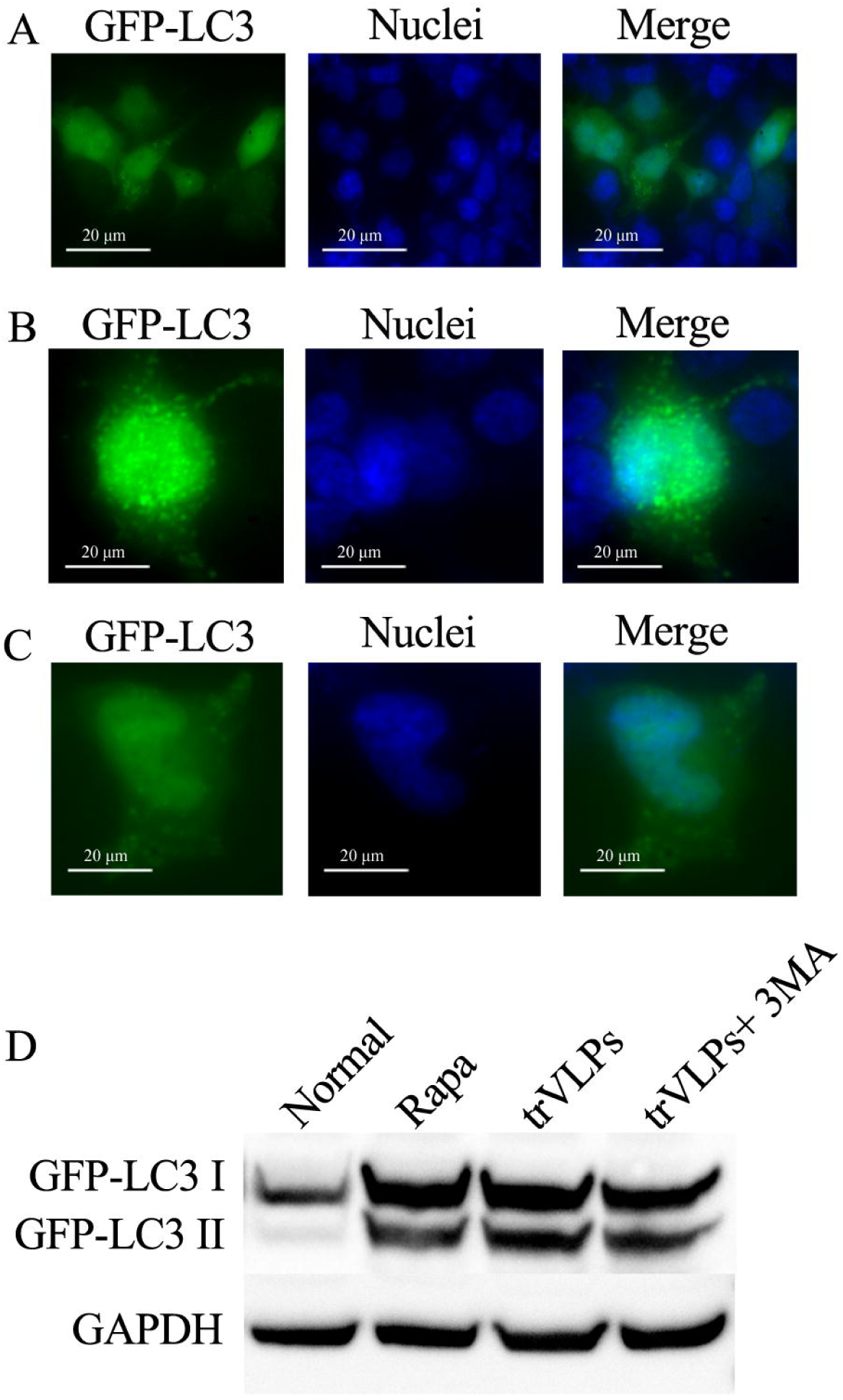
EBOV-trVLPs induce autophagy. (**A**) At 48 h post-trVLPs infection, autophagy was observed in 293T cells. GFP-LC3 punctate structures in the cells represent autophagosome numbers. **(B and C**) At a high magnification, rich GFP-LC3 punctate dots (≥ 5 dots/cell) are used to monitor autophagosome numbers in GFP-LC3-transfected cells. **(D)** Immunoblot analysis of autophagy. The conversion from GFP-LC3-I to GFPLC3-II indicated autophagy. The results indicated that the cells induced by rapamycin or infected by trVLPs were associated with significant GFP-LC3-II expression. Cells previously treated with 3-MA and then later infected with trVLPs also exhibited obvious GFP-LC3-II expression, which suggested that 3-MA could not effectively inhibit trVLPs-induced autophagy.

Thus, based on previous findings, including: 1) trVLPs could induce autophagy; 2) 3-MA could not significantly inhibit autophagy induced by trVLPs; 3) trVLPs GP, VP40 protein, and nucleic acids have been shown to interact with HSC70 and co-chaperones; and 4) GP and VP40 proteins have critical roles in the EBOV-trVLPs life cycle (e.g., virus entry, uncoating, fusion and budding processes), we presumed that trVLPs probably induced autophagy via the CMA pathway to benefit viral production. To prove this hypothesis, we assessed autophagy by the co-localization of LAMP1 and LC3 fluorescence, TEM observation, as well as an immunoblot analysis.

### EBOV-trVLPs induced autophagy via the CMA pathway

According to the protocol, 293T cells previously transfected with GFP-LC3 and RFP-LAMP1 were infected by EBOV-trVLPs. At 48 h post-infection, the co-localization of LC3 and LAMP1 was checked (Fig. 3A and B). Since LAMPs (e.g., LAMP1 and LAMP2) are lysosomal proteins and used as endosome and lysosome markers, RFP-LAMP1 can be found at the early stage of autophagolysosome formation during autophagy; moreover, GFP-LC3 was able to bind to autophagosomes and autolysosomes (Mizushima et al., 2001; Lee et al., 2008). Thus, the RFP-LAMP1-positive dots (red) represent lysosomes and autophagolysosomes; GFP-LC3-positive dots (green) indicate autophagosomes and autolysosomes; and the co-localization of RFP-LAMP1 and GFP-LC3-positive dots (yellow) indicate autolysosomes, which suggest the maturation of autophagy induced by the CMA pathway. Moreover, we investigated 293T cells infected with trVLPs for 48 h by TEM (Fig. 3C – H). In Fig. 3C, E, and G, autophagosomes (indicated by a single arrow) and autolysosomes (indicated by double arrows) are obviously visible in the cytoplasm. Fig. 3F clearly shows an autophagosome fused with a lysosome to form an autolysosome (indicated by double arrows).

**Figure 3.**
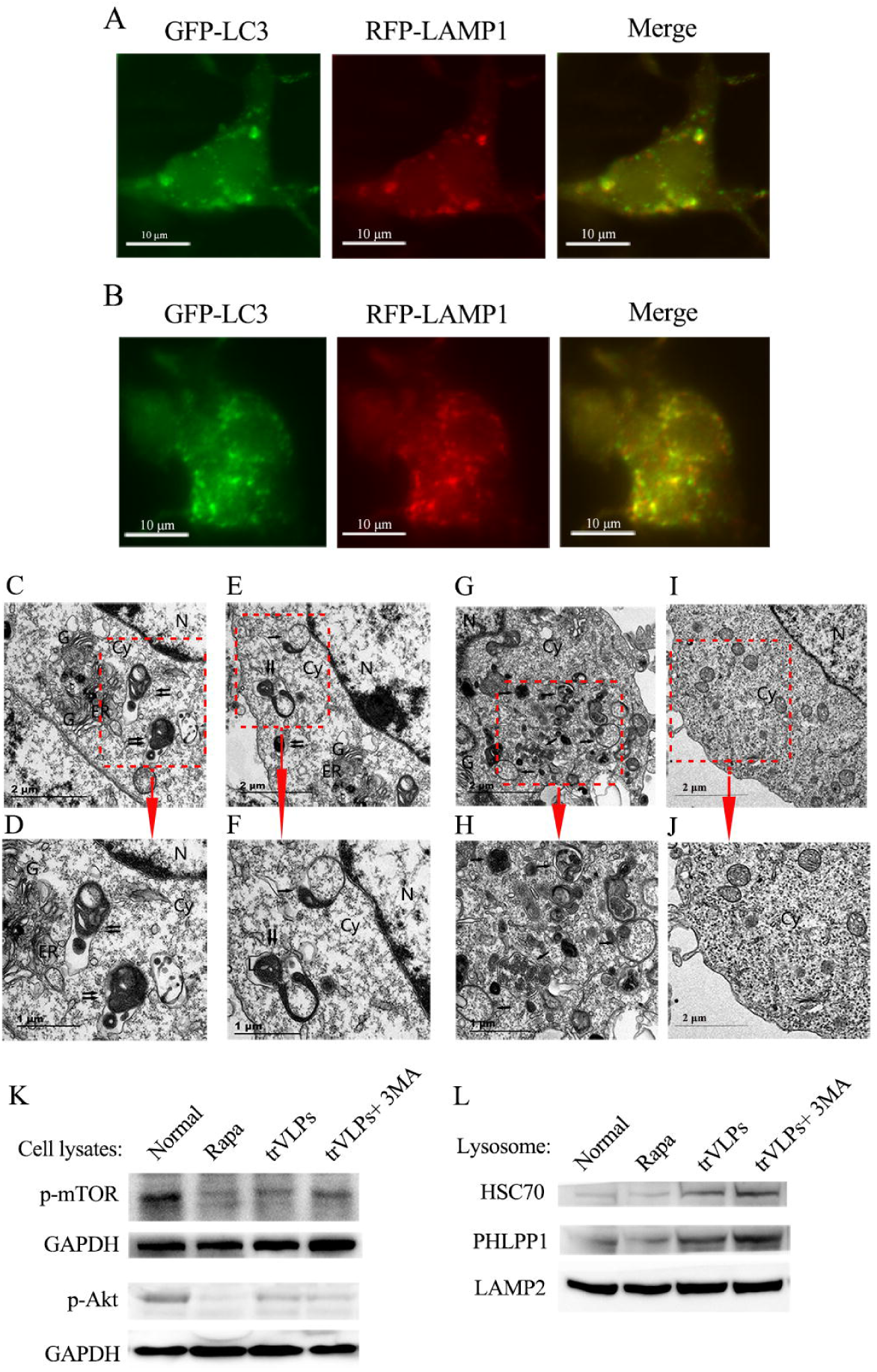
EBOV-trVLPs induced autophagy via the CMA pathway. **(A and B)** Co-localization of RFP-LAMP1- and GFP-LC3-positive dots (yellow) indicate autolysosomes. The 293T cells previously transfected with GFP-LC3 and RFP-LAMP1 were infected by EBOV-trVLPs for 48 h. RFP-LAMP1-positive dots (red) represent lysosomes and autophagolysosomes; GFP-LC3-positive dots (green) represent autophagosomes and autolysosomes. Thus, the co-localization of autophagosomes and autolysosomes is indicative of autolysosomes, which implies autophagy maturation induced by the CMA pathway. **(C** − **F)** Autophagosome and autolysosome morphology assessed by TEM. According to the protocol, 293T cells were infected with trVLPs for 48 h, then observed under TEM after sample preparation. In the images, autophagosomes (indicated by a single arrow) and autolysosomes (indicated by double arrows) are obviously visible in the cytoplasm. Panel F clearly shows an autophagosome fusing with a lysosome to form an autolysosome (indicated by double arrows). **(G and H)** 293T cells treated with rapamycin were used as a positive control. Autophagosomes (indicated by a single arrow) are richly expressed in the cytoplasm. **(I and J)** Normal 293T cells were treated as the negative control. **(K)** Immunoblot analysis of the level of cellular P-mTOR and P-Akt after infection with trVLPs. P-mTOR and P-Akt from the cell lysates were significantly lower in the cells treated with trVLPs or rapamycin or trVLPs plus 3-MA compared to the normal cells. Moreover, there were no distinct changes in P-mTOR or P-Akt expression between the groups treated with trVLPs and trVLPs plus 3-MA. **(L)** Immunoblot analysis of lysosomal HSC70 and PHLPP1 following infection with trVLPs. HSC70 and PHLPP1 were enriched in both the trVLPs and trVLPs plus 3-MA groups, but were thin in the rapamycin and normal groups. The lysosomal protein, LAMP2, was used as the internal control. Cy, cytoplasm; ER, endoplasmic reticulum; G, Golgi apparatus; N, nuclei.

Meanwhile, in order to confirm the expression of CMA related proteins, phosphorylated mTOR(P-mTOR) and Phosphorylated Akt (P-Akt) from cell lysates, HSPA1A and PHLPP1 from lysosomes were subjected to immunoblot analysis with related antibodies. The data indicate that P-mTOR was significantly lower in the cells treated with trVLPs or rapamycin compared to untreated cells. There were no distinct changes in P-mTOR expression in the cells treated with trVLPs or trVLPs plus 3-MA. P-Akt was associated with a highly similar trend with P-mTOR (Fig. 3I). In the lysosomes, HSPA1A was enriched in both trVLPs and trVLPs plus 3-MA treatment groups but appeared thin in the rapamycin and normal groups. PHLPP1 was substantially expressed in the trVLPs, rapamycin, and trVLPs plus 3-MA groups, compared with the normal cells (Fig. 3J). These results indicate that the Akt/mTOR/PHLPP1 axis is a common pathway for autophagy. In addition, lysosomes contribute to the CMA pathway, and the level of lysosomal HSPA1A is proportional to the level of CMA activity. To confirm that trVLPs could induce autophagy via chaperone or co-chaperone-related lysosomal activity, we knocked down HSC70, HSPA1A, and HSP90AB1, which revealed an interaction with trVLPs GP or VP40 proteins, an then examined its effect on CMA.

### TrVLPs-induced autophagy could be modulated by chaperone or co-chaperone-mediated pathways

According to the protocol, we knocked down HSC70, HSPA1A, and HSP90AB1 in 293T cells by siRNA, respectively, and infected the cells with trVLPs for 48 h. The knockdown efficiency of the siRNAs was verified in Fig. 4A. The expression of the GFP-LC3, P-mTOR, and P-Akt proteins were checked in the cell lysates. These data indicate that a knockdown of HSC70, HSPA1A, and HSP90AB1 could significantly inhibit the conversion of GFP-LC3-I to GFP-LC3-II but have little effect on P-mTOR and P-Akt (Fig. 4B). In addition, the knockdown of HSC70, HSPA1A, or HSP90AB1 could eliminate lysosomal HSC70, but had no effect on lysosomal PHLPP1 (Fig. 4C). The data also demonstrated that a knockdown of HSC70 and relevant co-chaperones could inhibit trVLPs-associated autophagy; however, there was no effect on the universal autophagy molecules, including mTOR, Akt, and PHLPP1. These findings indicate that trVLPs probably induced autophagy by several methods and was not limited to the CMA route.

**Figure 4.**
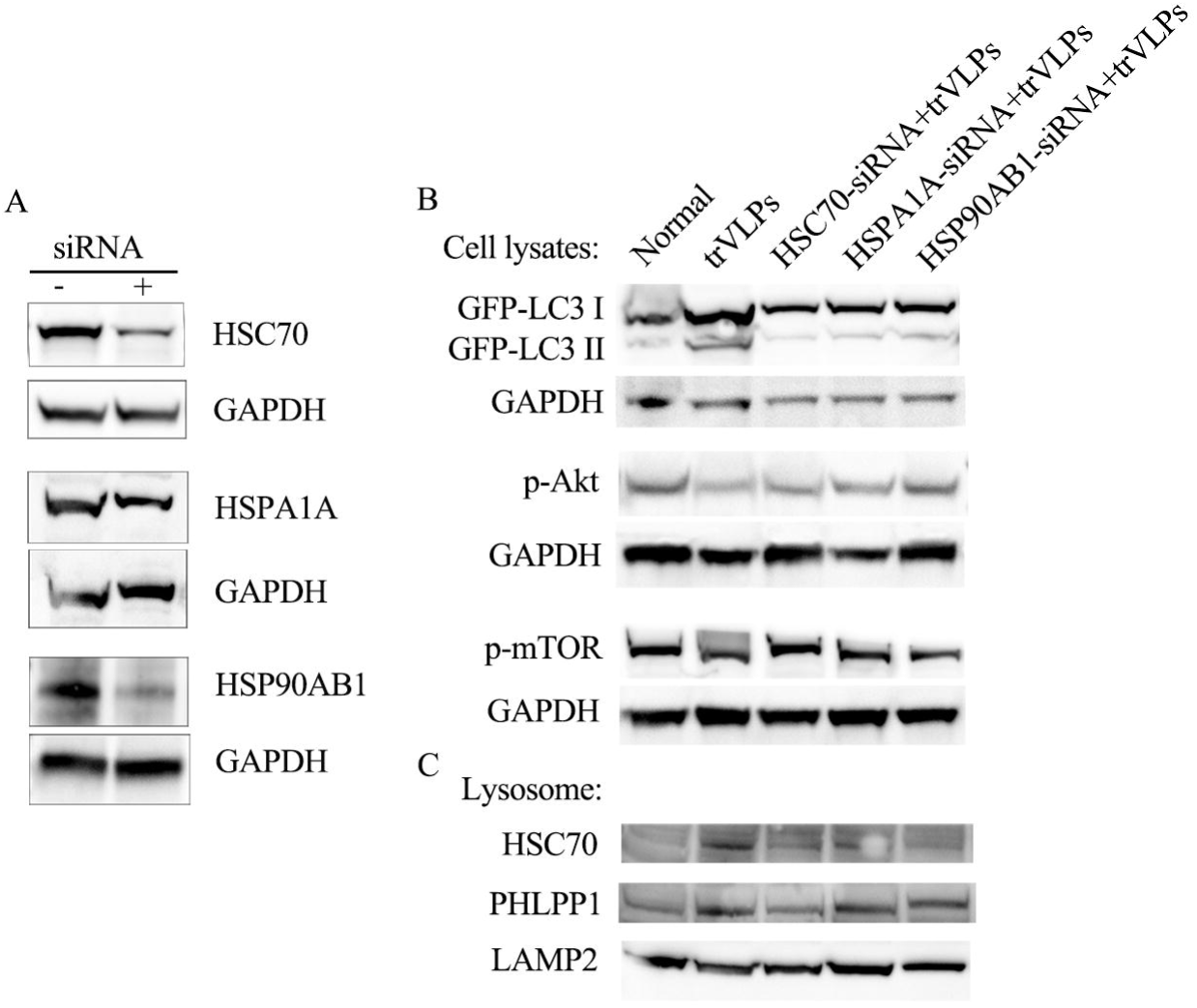
Downregulation of HSC70 and co-chaperones could affect trVLPs-induced CMA. **(A)** Western blot analysis of the target protein following siRNA transfection. HSC70, HSPA1A, and HSP90AB1 all exhibited a much lower expression after transfection with relevant siRNA compared with cells without siRNA interference. **(B)** An immunoblot analysis of cellular GFP-LC3, P-mTOR, and P-Akt following transfection of siRNA and infection with trVLPs. According to the protocol, the cells were previously transfected with siRNA and infected with trVLPs for 48 h. Knockdown of HSC70, HSPA1A, and HSP90AB1 could substantially inhibit the conversion of GFP-LC3-I to GFP-LC3-II, but had little effect on P-mTOR and P-Akt expression. **(C)** Immunoblot analysis of lysosomal HSC70 and PHLPP1 following siRNA transfection and trVLPs infection. Knockdown of HSC70, HSPA1A, and HSP90AB1 could eliminate lysosomal HSC70 expression, but had no effect on lysosomal PHLPP1. Lysosomal protein LAMP2 was used as an internal control.

## Discussion

To date, EBOV-induced cytopathy, including cellular necrosis, apoptosis, autophagy, and pyroptosis, urgently require further elucidation. Although EBOV has been shown to cause autophagy, detailed mechanisms remain unclear. In autophagy, Akt and mTOR represent major autophagy-inhibitory molecules, which plays an important role in autophagy signalling pathways and are regulated by various signals; P-mTOR and P-Akt are sensitive markers for assessing autophagy (Franke, 2008; Mirzaa et al., 2013).

In this study, we observed that EBOV-trVLPs infection significantly induced autophagy. Even pre-treatment with 3-MA, an inhibitor of class III PI3-K, could not inhibit trVLPs-induced autophagy. Since the PI3-K/Akt/mTOR pathway is a common autophagy pathway, this result indicated that trVLPs likely induced autophagy through other methods and are not limited to the PI3-K/Akt/mTOR pathway. In our previous study, we confirmed that the EBOV-trVLPs GP protein interacts with HSC70, HSPA1A, and HSP90AB1; moreover, viral nucleic acids and VP40 protein can both interact with HSPA1A and HSP90AB1 (Yu et al., 2018). Thus, we presumed that trVLPs most likely induced autophagy by chaperone or co-chaperone induced pathways. We assessed autophagy by the co-localization of RFP-LAMP1 and GFP-LC3 fluorescence and examined autophagy structures via TEM. Both the fluorescent images and TEM revealed autolysosome formation. Moreover, significantly low-expression of P-Akt and P-mTOR in the cell lysate, high PHLPP1 and HSC70 in the lysosome from trVLPs and trVLPs plus the 3-MA groups strongly suggest that trVLPs infection could induce autophagy via the CMA pathway, but is not limited to this pathway.

CMA is initiated with the binding of HSC70 to the substrates; however, a successful CMA process requires a series of coordinated events from other molecules, including co-chaperones and receptors (Tekirdag and Cuervo, 2018). It has been reported that a blockage of HSC70 co-chaperones present on the lysosomal surface could reduce substrate translocation, although the specific contribution of co-chaperones remains unclear (Agarraberes and Dice, 2001). Moreover, lysosomal HSP90, which localizes to both the cytosolic and luminal sides of the lysosome, facilitates LAMP2A stabilization as it transitions through different stages (Bandyopadhyay et al., 2008; Tekirdag and Cuervo, 2018).

In the siRNA interference tests, a knock down of HSC70, HSPA1A, and HSP90AB1 could inhibit the levels of autophagy and lysosomal HSC70, respectively; however, there was no effect on the P-Akt, P-mTOR, and PHLPP1 proteins. This data demonstrated that: 1) the down-regulation of HSC70 and co-chaperons could inhibit the CMA pathway but was negative to other autophagy pathways; 2) co-chaperons probably play an indispensable role in the CMA process; 3) low-levels of HSC70 and co-chaperons can possibly affect the viral protein-host protein interactions and further impair the trVLPs life cycle, which may indirectly affect the CMA.

There were limitations associated with our study. First, all of the tests were based on EBOV-trVLPs which may not reflect the physiology of live EBOV; thus, the results need to be confirmed under actual EBOV conditions. Second, except for CMA, we did not carefully check the other autophagy pathways that were probably induced by trVLPs. Finally, which of the trVLPs proteins primarily induced the CMA pathway and the detailed mechanisms remain unclear. In subsequent experiments, we will attempt to define the viral protein of trVLPs which induce CMA, clarify the mechanisms, and correlate these mechanisms with the trVLPs life cycle.

## Conclusion

In this study, we demonstrated that EBOV-trVLPs infection could induce autophagy by CMA but was not limited to this approach. In CMA, co-chaperons probably play an indispensable role. Moreover, it can be speculated that trVLPs GP or vp40 protein-host protein interactions probably affect autophagy, which might be related to the virus life cycle.

## Abbreviations

BAG-3: BCL2 associated athanogene-3;
BCA: Bicinchoninic acid;
BDBV: Bundibugyo virus;
CMA: Chaperone-mediated autophagy;
DiI: 1,1’-dioctadecyl-3,3,3’,3’-tetramethylindocarbocyanine perchlorate;
DMEM: Dulbecco’s modified Eagle’s medium;
EBOV: Ebola virus;
EVD: Ebola viral disease;
ER: Endoplasmic reticulum;
EM: Electron Microscopy;
FBS: Foetal calf serum;
HEK: Human embryonic kidney;
HSC70: Heat shock cognate protein of 70 kDa;
LAMP1: lysosome-associated membrane glycoprotein 1;
LC3: microtubule-associated protein light chain 3;
PAGE: Gel electrophoresis;
P-Akt: Phosphorylated Akt;
P-mTOR: phosphorylated mTOR;
RESTV: Reston virus;
RNP: Ribonucleoprotein;
SUDV: Sudan virus;
TAFV: Taï Forest virus;
TEM: Transmission electron microscopy.

## Acknowledgements

This work was supported by grants from the National Science and Technology Major Project for the Control and Prevention of Major Infectious Diseases in China (2018ZX10711001 and 2018ZX10102001).

## Author Contributions

DS Yu, SH Yao, WN Xi, LF Cheng, FM Liu, HB Hu, XY Lu and HP Yao performed the experiments. HP Yao, DS Yu performed the statistical analysis. HP Yao, DS Yu, NP Wu and SL Sun designed the study and drafted the manuscript. All authors participated in writing the manuscript.

