## Supplementary material for "HSC70, HSPA1A, and HSP90AB1 Facilitate Ebola Virus trVLPs to Induce Autophagy": Supplementary Figure 1.pdf

Supplementary Figure 1. Evaluation of cytotoxity of siRNAs by 293T cells nuclei counting in one vision of microscope (40×, 0.24mm$^2$). (A) Evaluation of cytotoxic effects of HSC70 siRNA. 293T cells (1×10$^4$ cells/well in 48-wells plate) were transfected with HSC70 siRNA (75 nM) for 48 h, then nuclei were strained with DAPI and counted, the siRNA was classified as hypotoxicity as 75 or more nuclei in one vision of microscope (40×, 0.24mm$^2$). (B) and (C) similar assessments were done for HSPA1A and HSP90AB1 siRNAs, respectively.

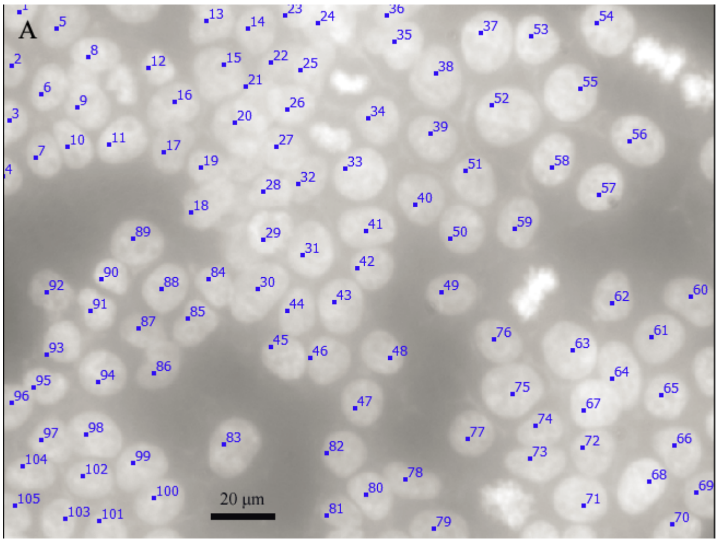


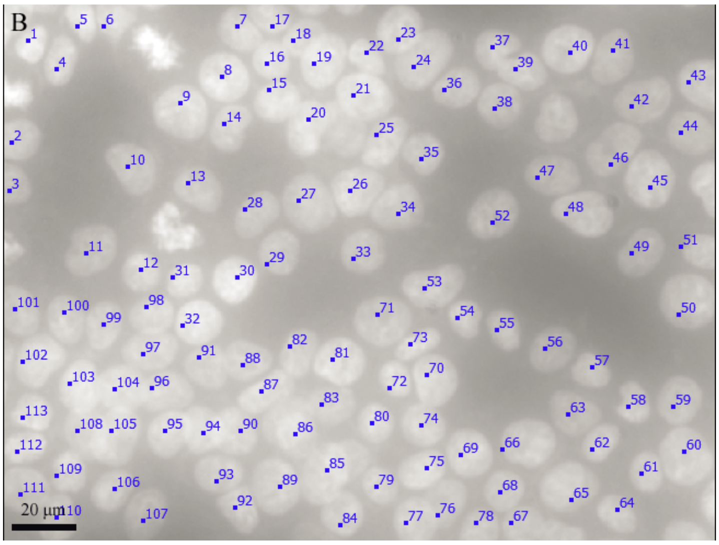

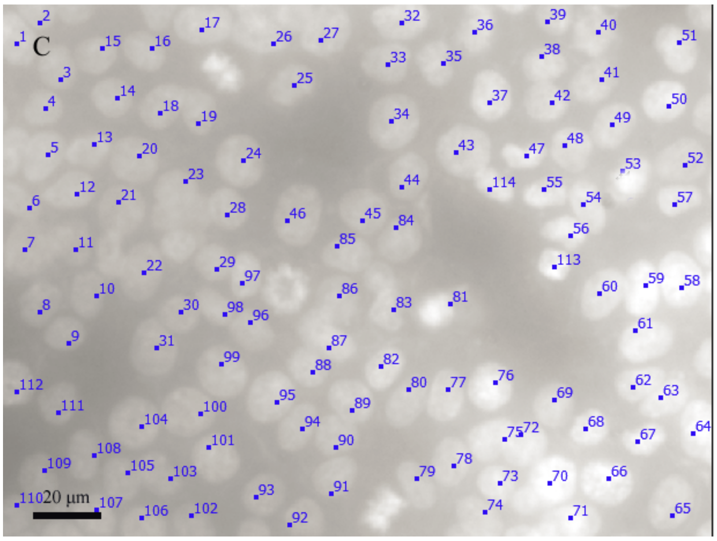
C
20 µm
