## Supplementary material for "HSC70, HSPA1A, and HSP90AB1 Facilitate Ebola Virus trVLPs to Induce Autophagy": Supplementary table 1.pdf

Supplementary table 1. TCID50 of trVLPs calculated by Reed-Muench method and conversion of MOI values.

| Dilution of trVLPs | Counts of CPE wells | Counts without CPE wells | Accumulation of CPE wells | Accumulation without CPE wells | Percentage of CPE wells (%) |
|---|---|---|---|---|---|
| 10-1 | 7 | 1 | 18 | 1 | 94.7（18/19） |
| 10-2 | 5 | 3 | 11 | 4 | 73.3（11/15） |
| 10-3 | 3 | 5 | 6 | 9 | 40.0（6/15） |
| 10-4 | 2 | 6 | 3 | 15 | 16.7（3/18） |
| 10-5 | 1 | 7 | 1 | 22 | 4.3（1/23） |
| 10-6 | 0 | 8 | 0 | 30 | 0（0/31） |
| 10-7 | 0 | 8 | 0 | | |

Cytopathic effect= (73.3-50)/ (73.3-40) =23.3/ 33.3= 0.70

lgTCID50=0.70× (-2) + (-2) = -3.4

TCID50=10E-3.4/ 0.1ml

MOI= TCID50×0.7/cell numbers =TCID50×0.7/10000 cells= 0.20
