## Supplementary material for "HSC70, HSPA1A, and HSP90AB1 Facilitate Ebola Virus trVLPs to Induce Autophagy": Supplementary table 2.pdf

Supplementary table 2. Information of siRNAs chosen for RNA interference experiments.

| **NO.** | **siRNAs (product name)** | **Product Id** | **siRNA Target Sequence** | **Ct （primer：gene）** | **Ct（primer：18s）** | **ΔCt** | **-ΔΔCt** | **$2^{-\Delta\Delta Ct}$** |
|---|---|---|---|---|---|---|---|---|
| 1 | negative control | | | 27.86 | 15.81 | 12.05 | | |
| | Hs_HSC70 | SI04213370 | GAGGTTGATTAAGCCAACCAA | 28.85 | 14.89 | 13.96 | -1.91 | 0.2660925 |
| 2 | negative control | | | 36.4 | 13.67 | 22.73 | | |
| | Hs_HSPA1A | SI04364136 | TCCGGTTTCTACATGCAGAGA | 37.42 | 14.78 | 22.64 | 0.09 | 1.0643702 |
| 3 | negative control | | | 21.6 | 13.27 | 8.33 | | |
| | Hs_HSP90AB1 | SI02780561 | CAAGAATGATAAGGCAGTTAA | 22.8 | 14.18 | 8.62 | -0.29 | 0.8179021 |
