## Supplementary figures and images for "HSC70, HSPA1A, and HSP90AB1 Facilitate Ebola Virus trVLPs to Induce Autophagy"

### Original pictures of Blot.pdf

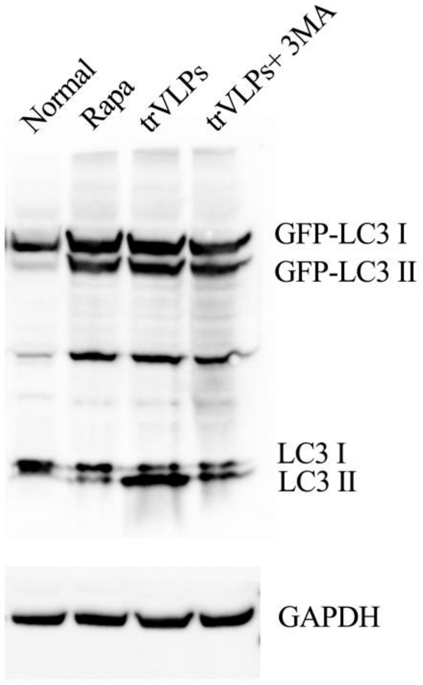

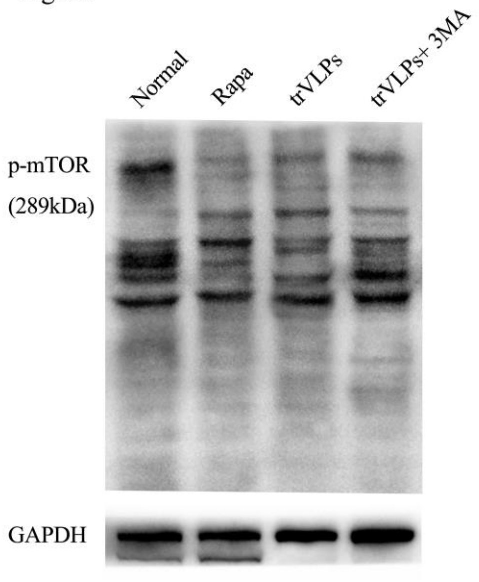

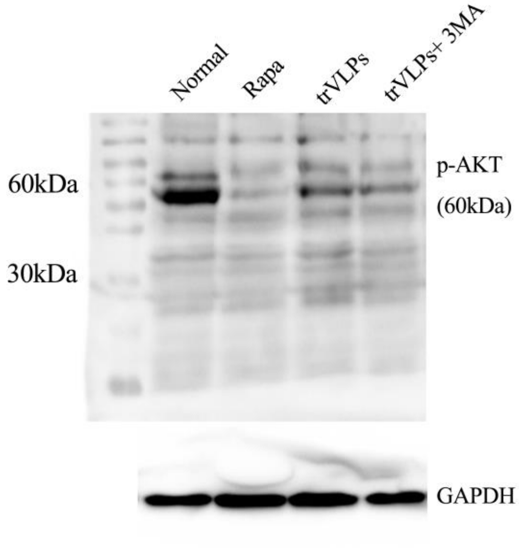

Fig. 3L

Lysosome:

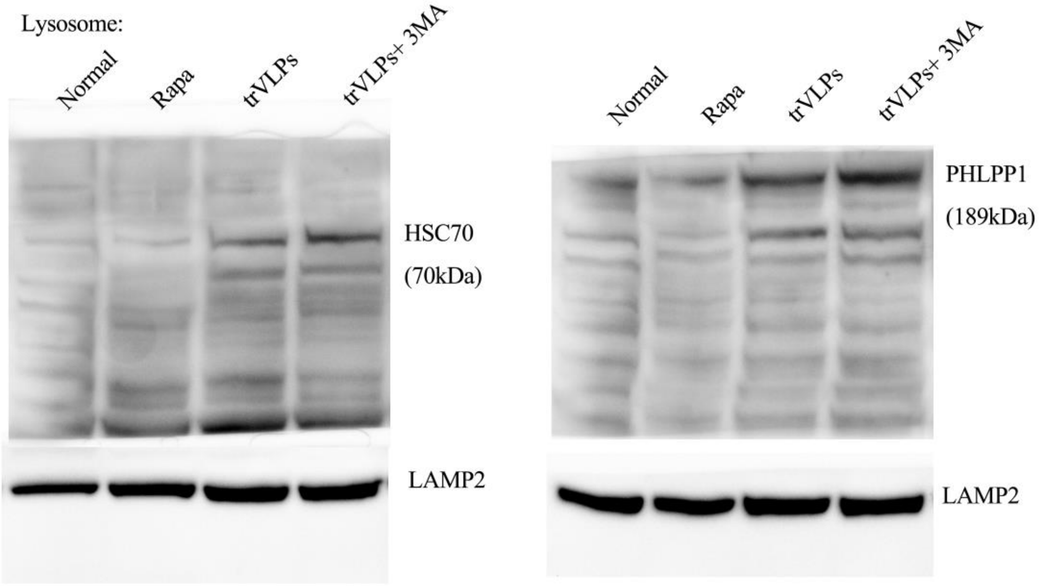


Fig. 4A

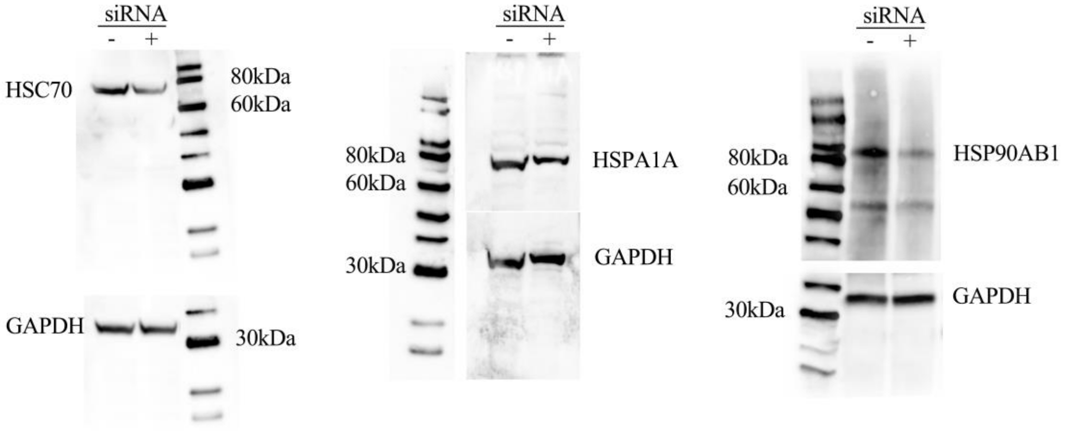

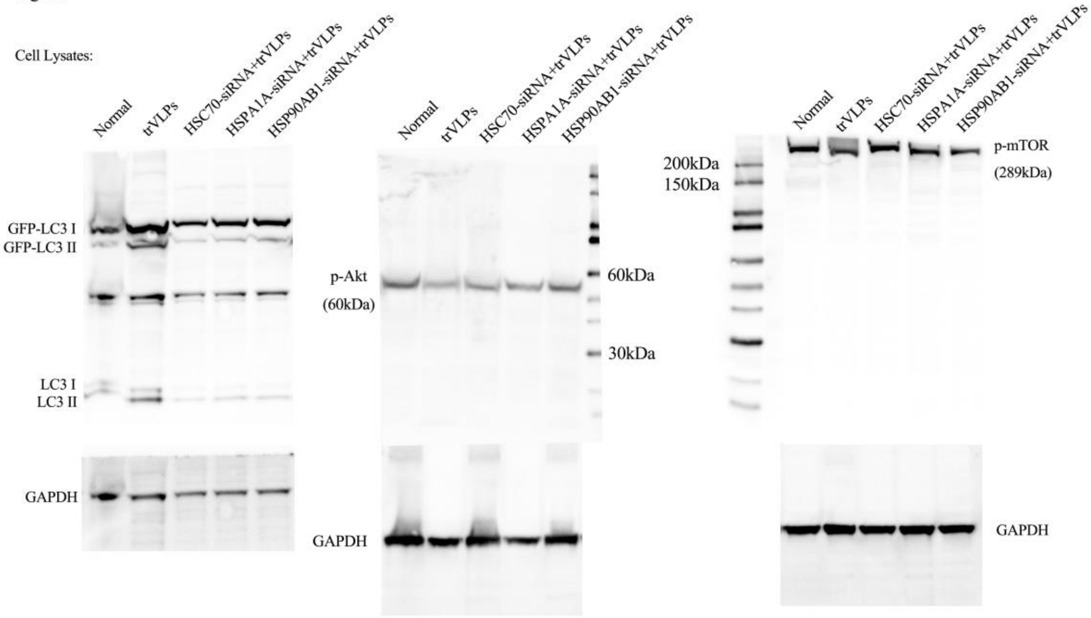

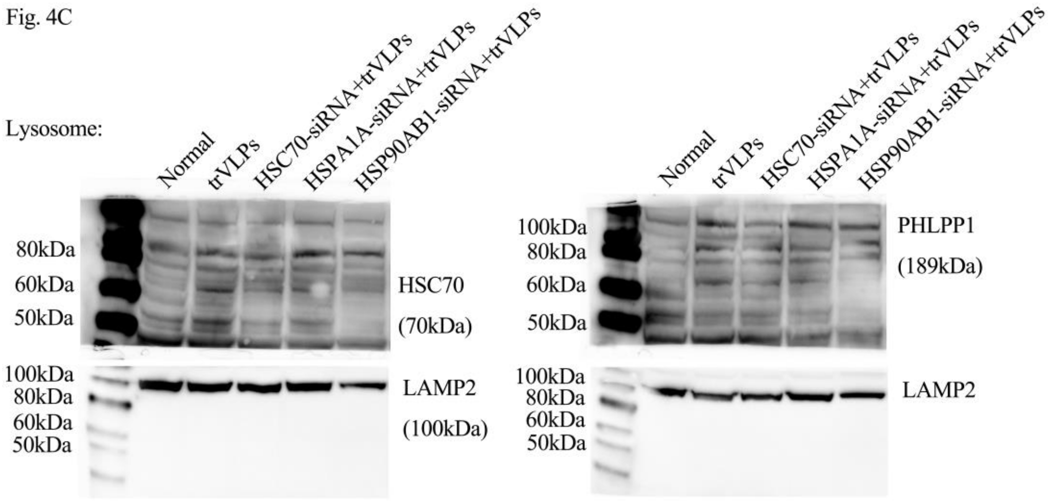
